# Impacts of 203/204: RG>KR mutation in the N protein of SARS-CoV-2

**DOI:** 10.1101/2021.01.14.426726

**Authors:** Majid Vahed, Tess M Calcagno, Elena Quinonez, Mehdi Mirsaeidi

## Abstract

We present a structure-based model of phosphorylation-dependent binding and sequestration of SARS-CoV-2 nucleocapsid protein and the impact of two consecutive amino acid changes R203K and G204R. Additionally, we studied how mutant strains affect HLA-specific antigen presentation and correlated these findings with HLA allelic population frequencies. We discovered RG>KR mutated SARS-CoV-2 expands the ability for differential expression of the N protein epitope on Major Histocompatibility Complexes (MHC) of varying Human Leukocyte Antigen (HLA) origin. The N protein LKR region K203, R204 of wild type (SARS-CoVs) and (SARS-CoV-2) observed HLA-A*30:01 and HLA-A*30:21, but mutant SARS-CoV-2 observed HLA-A*31:01 and HLA-A*68:01. Expression of HLA-A genotypes associated with the mutant strain occurred more frequently in all populations studied.

**Importance:** The novel coronavirus known as SARS-CoV-2 causes a disease renowned as 2019-nCoV (or COVID-19). HLA allele frequencies worldwide could positively correlate with the severity of coronavirus cases and a high number of deaths.

## 1. Introduction

A major outbreak of a novel coronavirus isolated in china called SARS-CoV2 leads to the global declaration of a worldwide pandemic (1). The novel coronavirus known as SARS-CoV2 causes a disease renowned as 2019-nCoV (or COVID-19) (1-3). Sequencing of the viral genome and characterization of immunogenic viral proteins has been crucial for understanding the virus and creating targeted vaccinations and treatments. Additionally, wide variation in the clinical presentation of COVID-19 ranging from an asymptomatic presentation to disabling multi-organ failure has led to the study of differential host-pathogen interaction specifically relating to population-based Human Leukocyte antigen (HLA) allele frequencies (4, 5).

SARS-CoV2 is an RNA virus and structurally contains 16 open reading frames (ORFs). ORF1a express 11 nonstructural proteins (Nsps) from Nsp1 to Nsp11, with the genes of ORF1b expressing proteins from Nsp12 to Nsp16 (6, 7). Major structural proteins including Spike (S), envelope (E), membrane (M) and nucleocapsid (N) proteins are encoded by other ORFs. The N proteins of SARS-CoV are highly basic structural proteins localized in the cytoplasm and the nucleolus of Trichoplusia ni BT1 Tn 5B1-4 cells (8). Previous studies have indicated the N proteins of other coronaviruses are extensively phosphorylated and bound to viral RNA to form a helical ribonucleoprotein (RNP) that comprises the viral core structure (9). Recently, mutations in N segment of SARS-CoV2 have been reported (10). Two replacements in positions R203K and G204R of N proteins have been found in several countries, but their potential effects in the protein structure have not been discussed.

We present a structure-based model of phosphorylation-dependent binding and sequestration of SARS-CoV-2 nucleocapsid protein and the impact of two consecutive amino acid changes R203K and G204R. Additionally, we studied how mutant strains affect HLA-specific antigen presentation and correlated these findings with HLA allelic population frequencies.

## 2. Results

### 2.1. Sequence Alignments and Clustering of LKR of CoVs N-protein

ClustalW multiple sequence alignment was employed to align the LKR (68 nucleotides long) of CoV N protein aligned for bat/pangolin models and human models SARS-COVs/MERS-COV. Notably, The Gly at position 204 is conserved among the closely related coronaviruses Bat coronavirus pangolin, SARS-CoV, and SARS-CoV-2, but variable for SARS-CoV2n (fig. 1(a)). The clustering trees of coronavirus are displayed in figure 1 (b). Sequence alignments suggested that other coronavirus N proteins might share the same structural organization based on intrinsic disorder predictor profiles and secondary structure predictions (Fig. 2).

**Figure 1.**
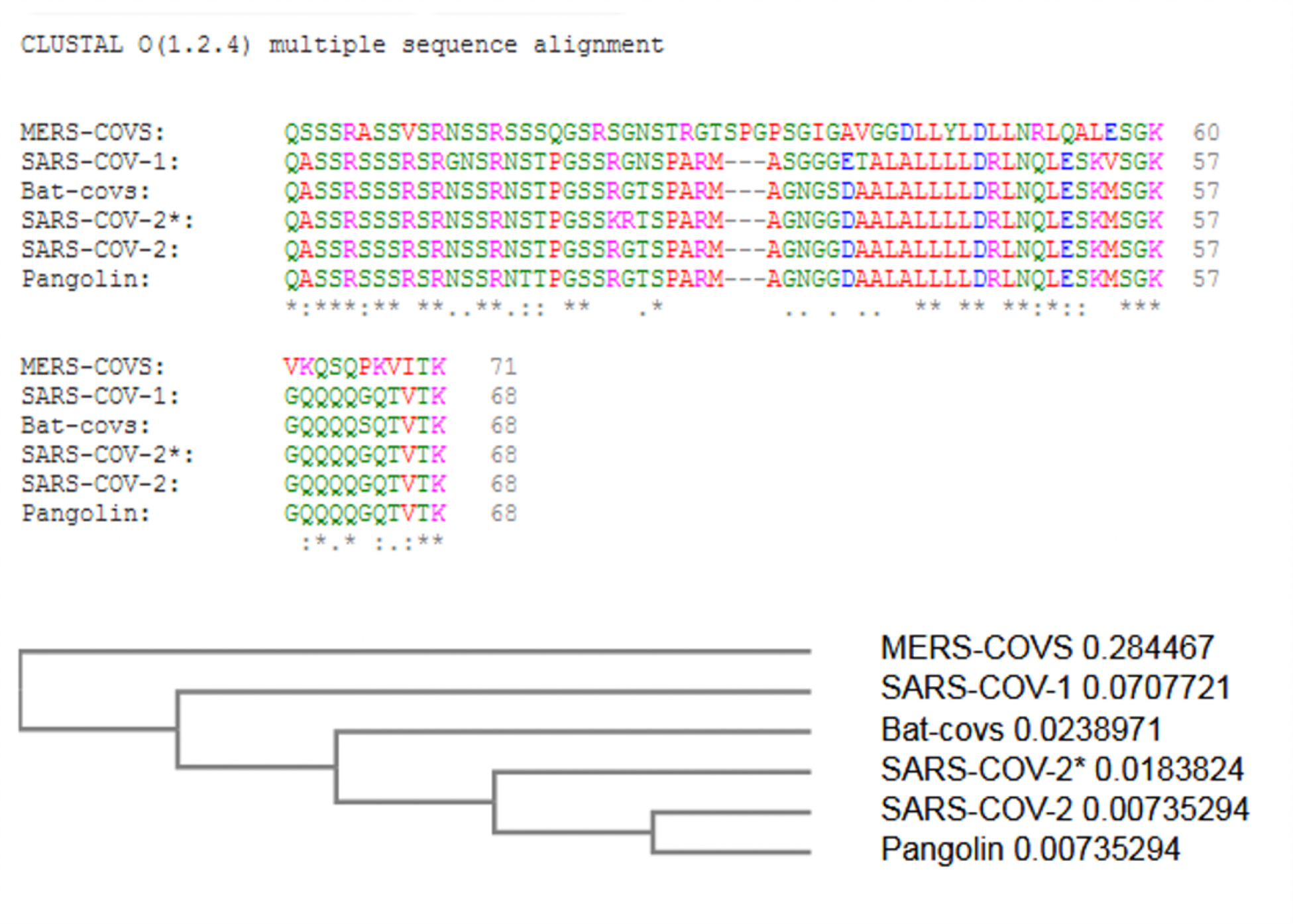
Coronavirus N-protein LKR sequence domains. (a) Alignment for bat coronavirus pangolin, SARS-CoV, SARS-CoV-2 and SARS-CoV2n each genotype for sequence representation. Columns with changes for nucleotide positions have been color-coded. (b) ClustalW multiple sequence alignment trees display of coronavirus. (*)The asterisk indicates the mutation type.

**Figure 2.**
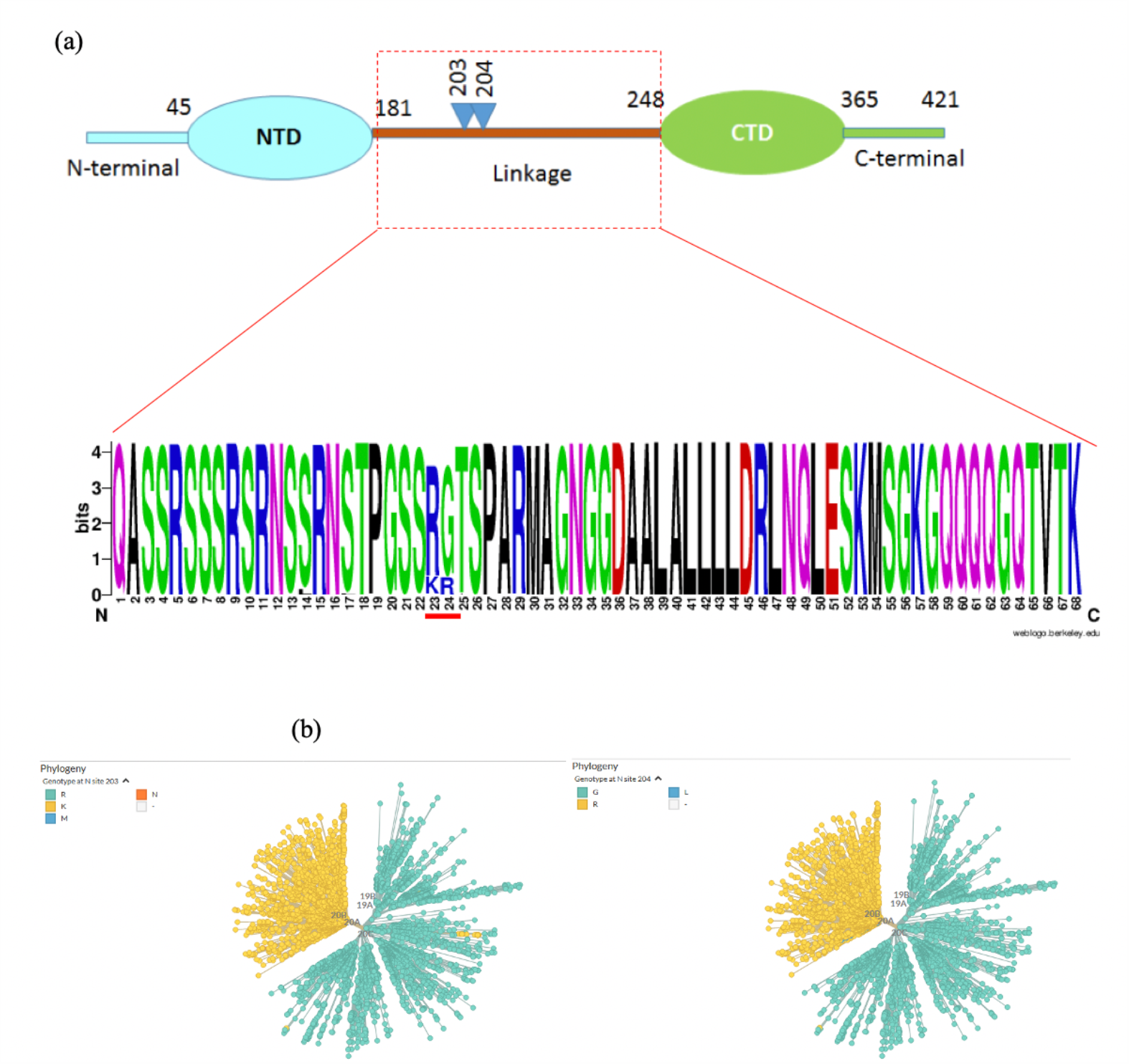
(a) Schematic representation of the N protein LKR domain, (b) Multiple sequence alignment logos of selected N protein LKR domain, a.a. letter heights indicate their frequency at the position 203/204 in SARS-COV-2. (c) The SARS-CoV-2 N protein conserved mutations at the position 203/204 in a phylogenetic graph obtained from Nextstraindatabase. Picture captured date: 9/25/2020.

### 2.2. Calculation of N-protein LKR Residue Energies

A plot of N-proteins residues indicates the local model quality by plotting knowledge-based energies as a function of an amino acid sequence position. A detailed energy calculation of wild and mutant types revealed residues 203/204 are located in the highest energy level area (Fig. 3). The mutant type showed slightly high free energy at residues 203/204 a.a. KR compared to the wild type; suggesting enhanced structural flexibility and increased the tendency for the formation of a coil or a bend in the secondary structure. The relative orientation of NTD and CTD, as well as the conformations of the disordered regions (N-arm, LKR, and C-tail), are drawn randomly to reflect the dynamic nature of the N protein (Fig. 3)

**Figure 3.**
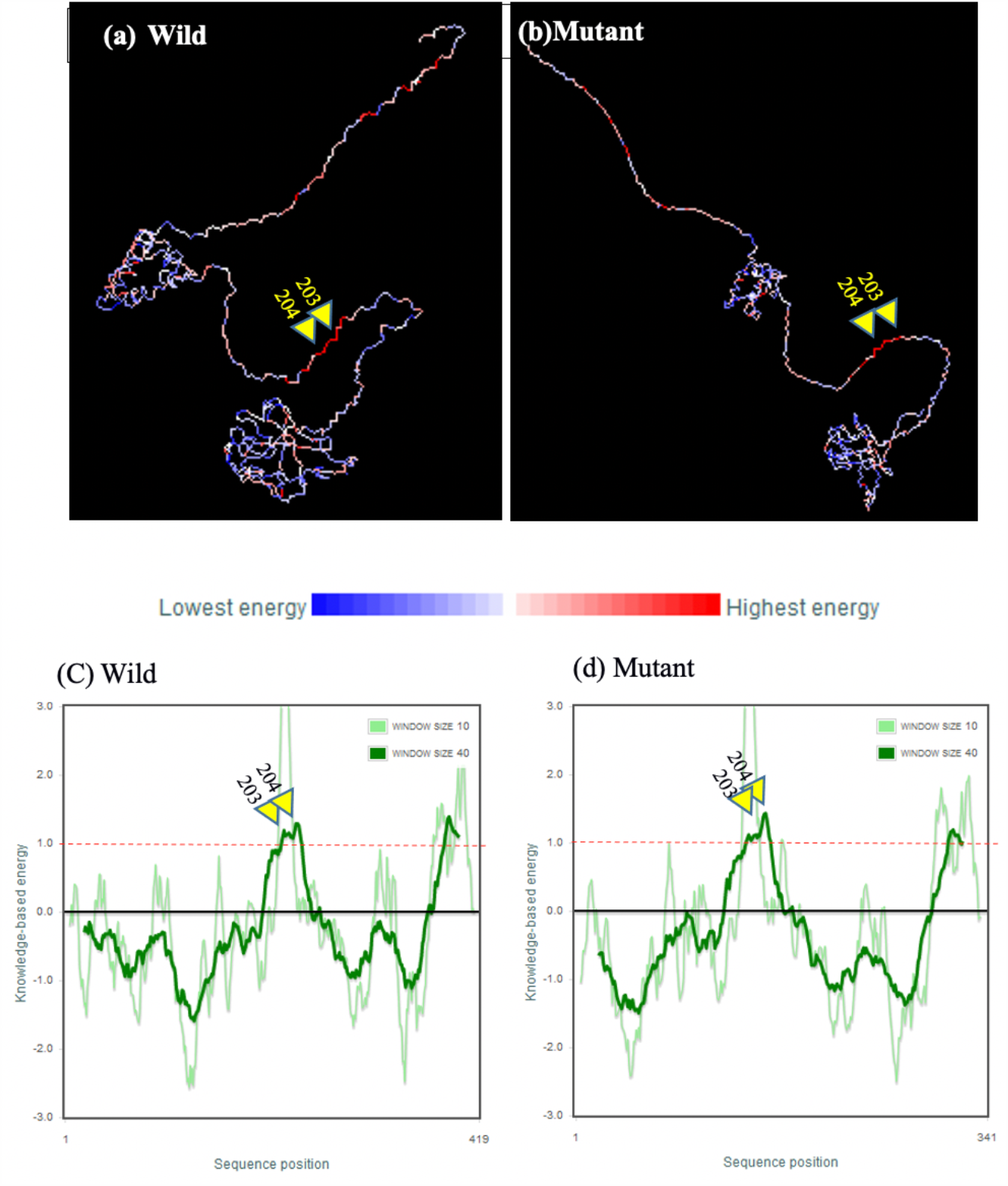
Three structure dimensional depicted of the based on the energy values where “blue” represents lowest energy and “red” represents highest energy in the N-protein (a) SARS-CoV-2 wild (b) SARS-CoV-2 mutant. (c) ProSa energy plots of N-protein wild type a.a. 203/204 RG (d) mutant type a.a. 203/204 KR.

### 2.3. Calculation of Putative Phosphorylation Sites

We also analyzed predictable phosphorylation sites of N-protein by employing the NetPhos 3.1 server (http://www.cbs.dtu.dk/services/NetPhos/, accessed on 14 Sep 2020). The linker region of SARS-CoV N-protein (LKR) contains a Ser/Arg-rich region with a high number of putative phosphorylation sites (a.a. 172–206). The sites of contiguous amino acid changes of 203R>K and 204G>R are located in the SR-rich region which is known to be intrinsically disordered. We predicted a nonspecific kinase phosphorylation site at Ser202 and a specific CDK5, RSK, and GSK3 phosphorylation sites at Ser206, all of which are close to the RG>KR mutation (Fig. 4). When Ser202/206 and Thr205 are phosphorylated, charge neutralization of the nearby positively charged sidechains likely takes place. The G204R mutation decreases the conformational entropy of neutralization by increasing the positive charge in the vicinity of negatively charged P-Ser202/205 phosphate groups (Fig. 5).

**Figure 4.**
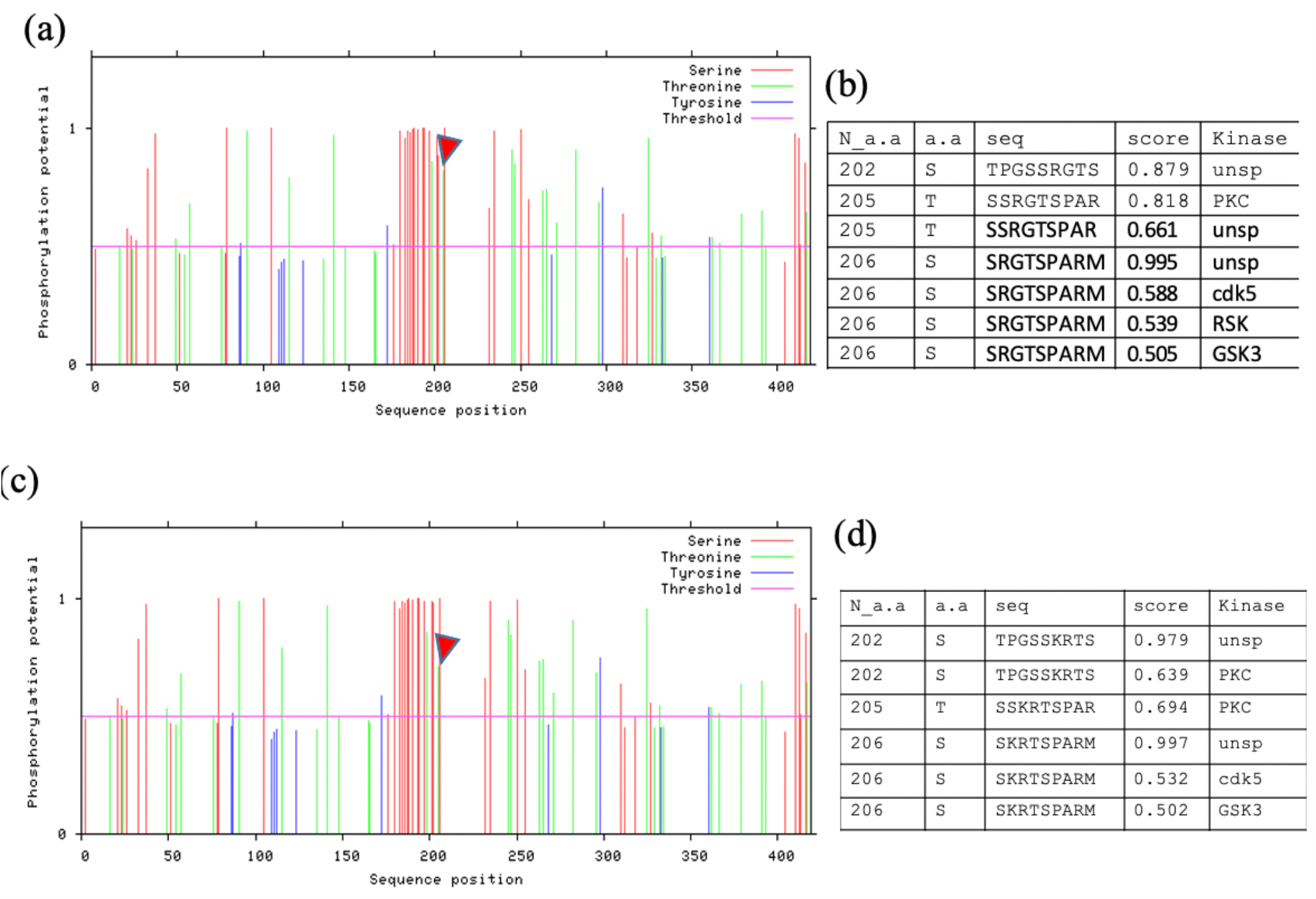
ProSa Phosphorylation plots of N-protein. (a) Phosphorylation sites predicted in the local LKR SARS-CoV-2 wild (b) phosphorylation score and substrates of kinases SARS-CoV-2 wild (c) Phosphorylation sites predicted in the local LKR SARS-CoV-2 mutant.(d) phosphorylation score and substrates of kinases SARS-CoV-2 mutant.

**Figure 5.**
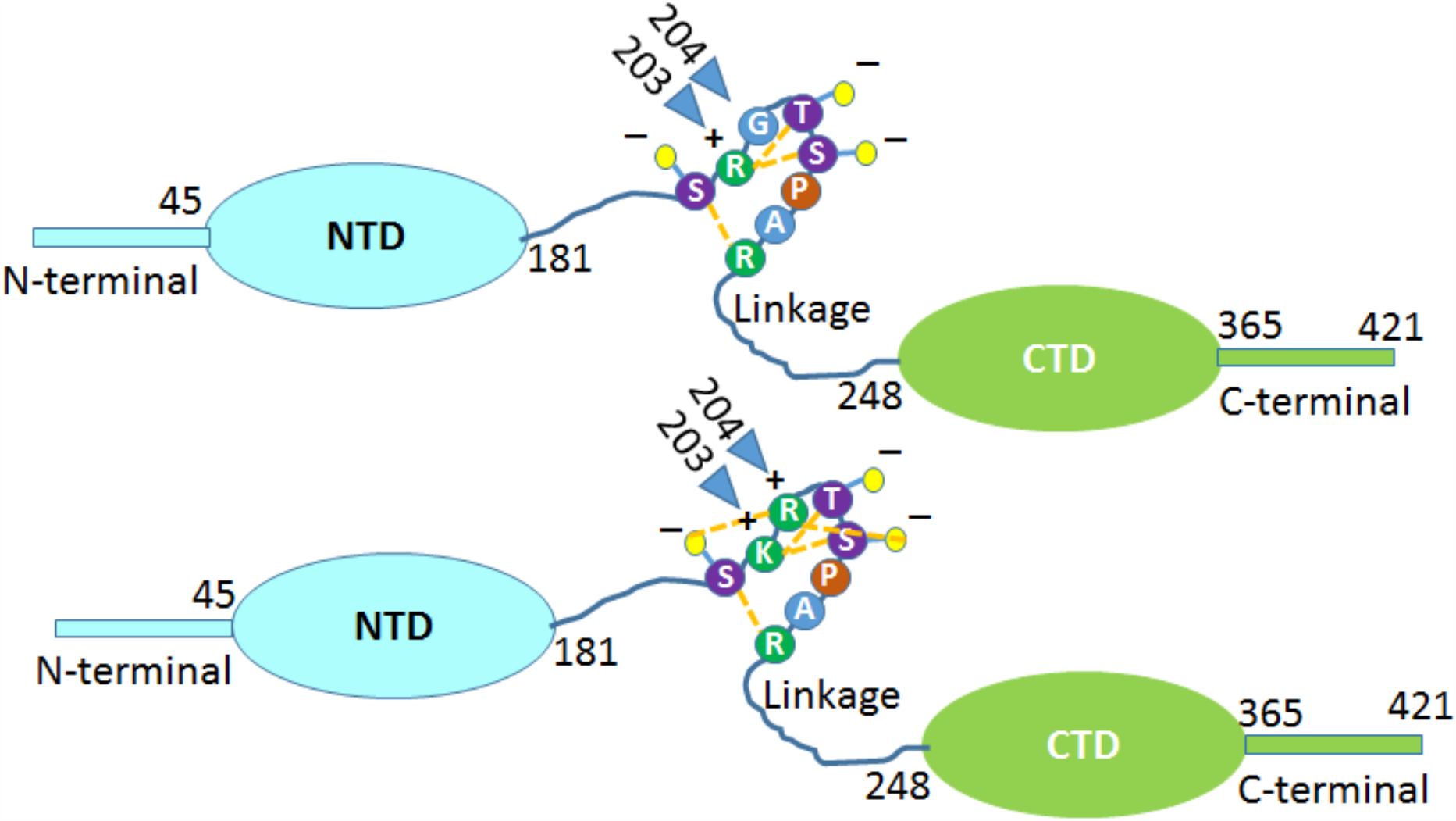
Schematic representation of the mutation and possibility electrostatic interactions between positively charged Arg/Lys and the negatively charged P-Ser/Thr as shown yellow dotted lines. (a) the wild type SARS-CoV-2 N-protein with Arg and Gly at position 203 and 204, respectively (b) the variant SARS-CoV-2 N-protein with mutations at position 203 and 204 (203R>K, 204G>R). These residues are colour-coded based on their Charges (Green: positively charged, Purple: negatively charged P-Ser/Thr. The phosphate groups on Ser/Thr residues at position 202, 205 and 206 are denoted as red circles on yellow sticks

### 2.4 Protein Localization

Subcellular localization of the N-protein was predicted using the DeepLoc 1.0 neural network algorithm. The resultant values of 0.861 and 0.913 obtained for wild and mutant N protein respectively suggest the protein is predominantly found in the nucleus (Fig. 6 a,b). The N-protein position prediction graph (Fig.6,c,d) confirmed a long peak in mutation areas K203 and R204, defined based on SARS-CoV-2 data.

**Figure 6.**
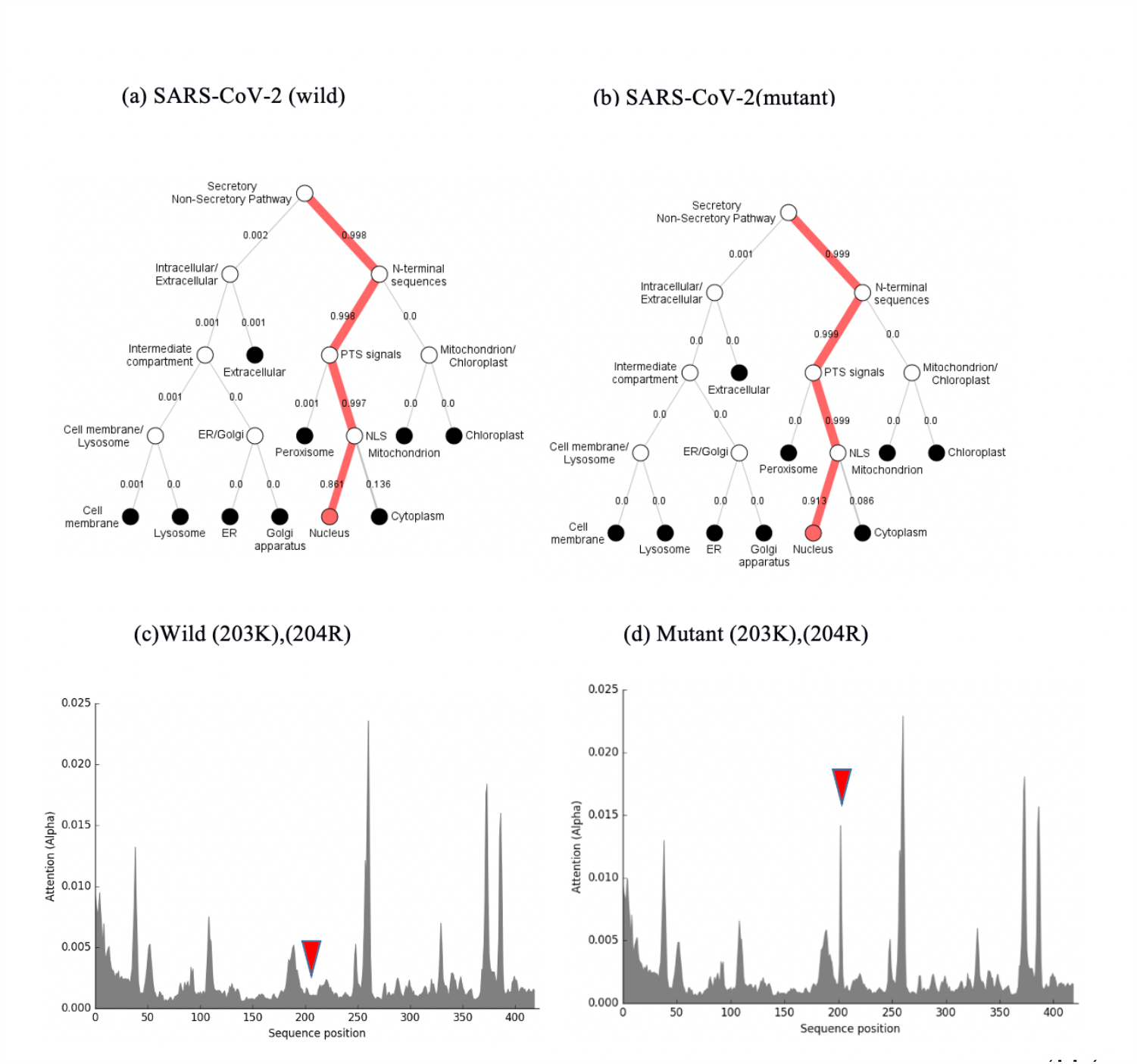
Hierarchical tree-predicted subcellular localizations of N-protein using neural networks algorithm. (a) Wild (203R),(204G). (b) Mutant (203K),(204R). (c) N-protein residue position prediction Wild (203K),(204R). (d) N-protein residue position prediction Mutant (203K), (204R)

### 2.5 Identification of N-protein B-cell epitopes

We used the Immune Epitope Database (IEDB) to determine linear B-cell hosted epitopes utilizing the incorporated Chou & Fasman Beta-Turn prediction module threshold 1.07. We supplied the FASTA sequence of the targeted protein as an input assuming all default parameters. LKR (a.a.170-206) region of N-protein shown significant antigenic epitopes with potential binding to B lymphocytes cells (Fig. 7). IEDB software also predicted epitopes based on N-protein conformation and residue exposure, and independently graphed a broad peak in mutation areas K203 and R204, the likely epitope regions defined based on SARS-CoV-2 data (Fig. S2).

**Figure. 6.**
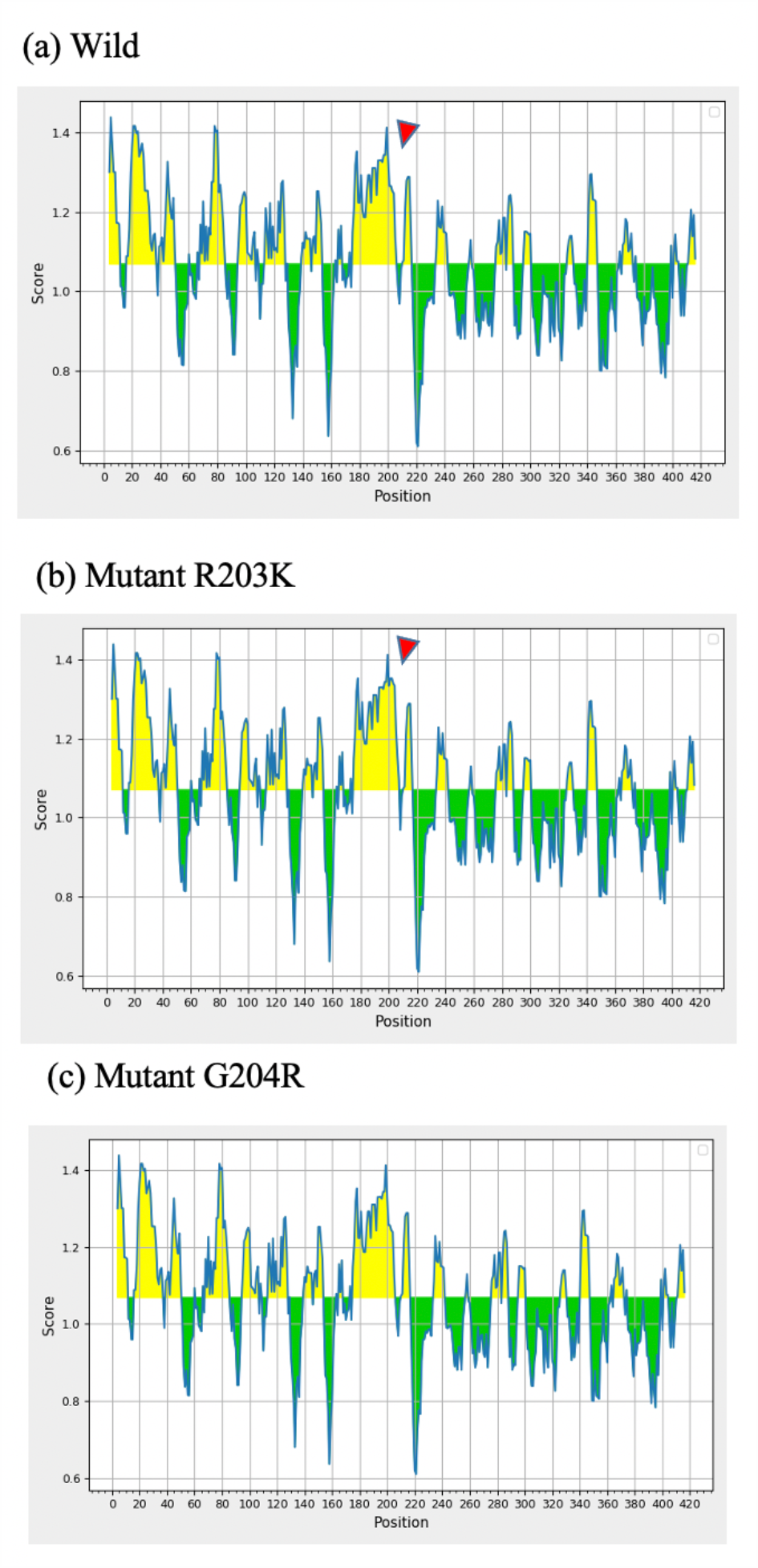
Graphical representation of prediction linear B-cell epitopes within the N-protein of (a) SARS-COV-2 wild (b) SARS-COV-2 R203K mutant (c) G203R by IEDB scales. Arrow depicts the residue 204.

### 2.6 Identification of T-cell epitopes

We found that MHC polymorphism typically results in differential MHC epitope recognition within the N-protein LKR K203 R204 region of wild type (SARS-CoV-2) as compared to mutant (SARS-CoV2). Wild type (SARS-CoV-2) observed HLA-A*30:01 and HLA 31:01 predictions, but mutant SARS-CoV-2 observed HLA-A*31:01 and HLA-A*68:01 predictions (Table 1,2). The frequency of HLA class I representation within South American, Japanese, and Iranian populations was recorded for both wild type and mutant strains. HLA class I molecules associated with mutant strains occurred more frequently in all three population groups, with the most significant increase seen in the South American population (18.38% in wild type versus 28.25% in the mutant).

**Table 1:**
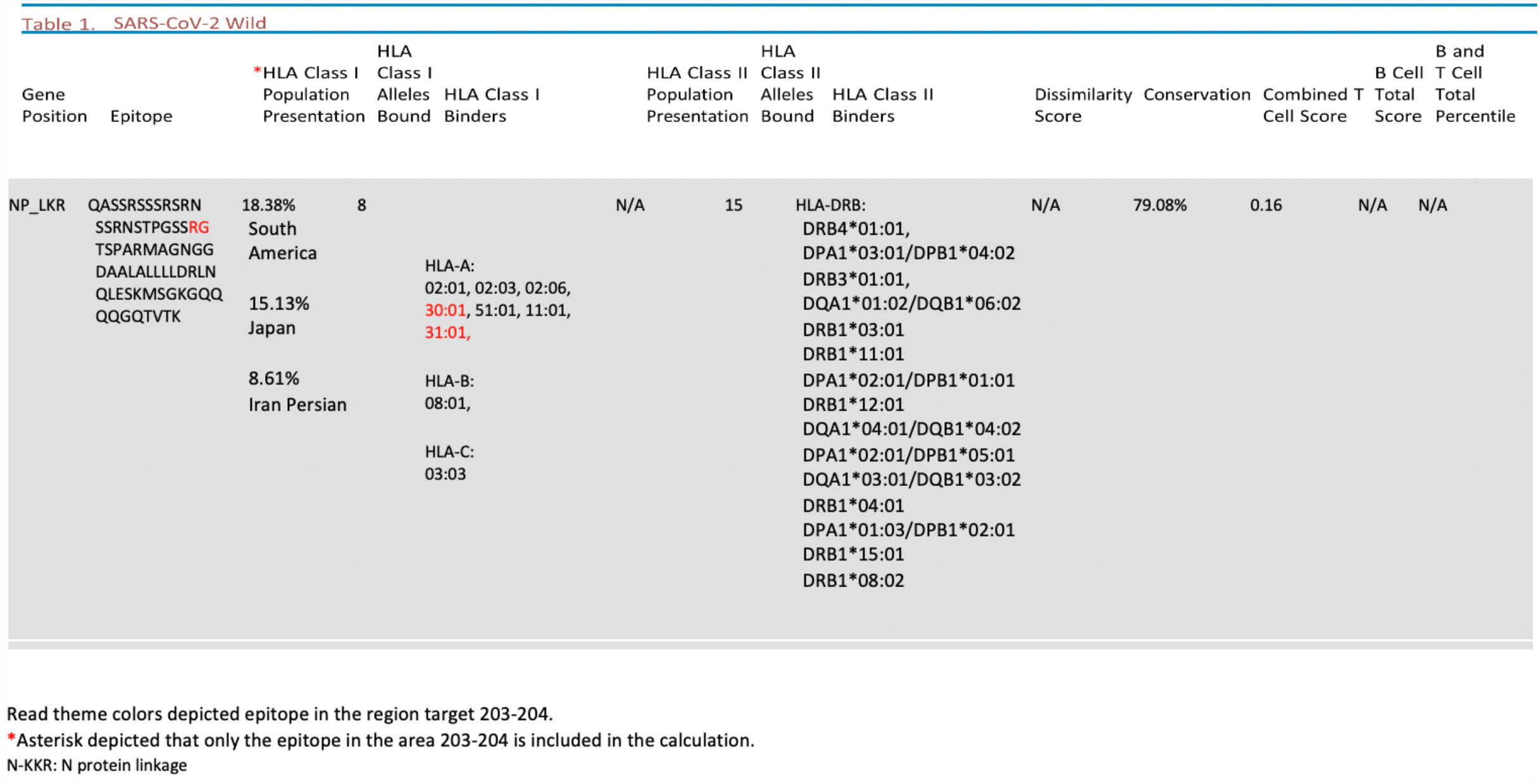
SARS-CoV-2 wild type N protein T cell HLA epitope predictions at gene position NP_LKR. HLA-class I binders depicted in red included only the epitope in the target region 203-204 in the calculation. Frequency of HLA class I representation within South American, Japanese, and Iranian populations is recorded as a frequency.

**Table 2:**
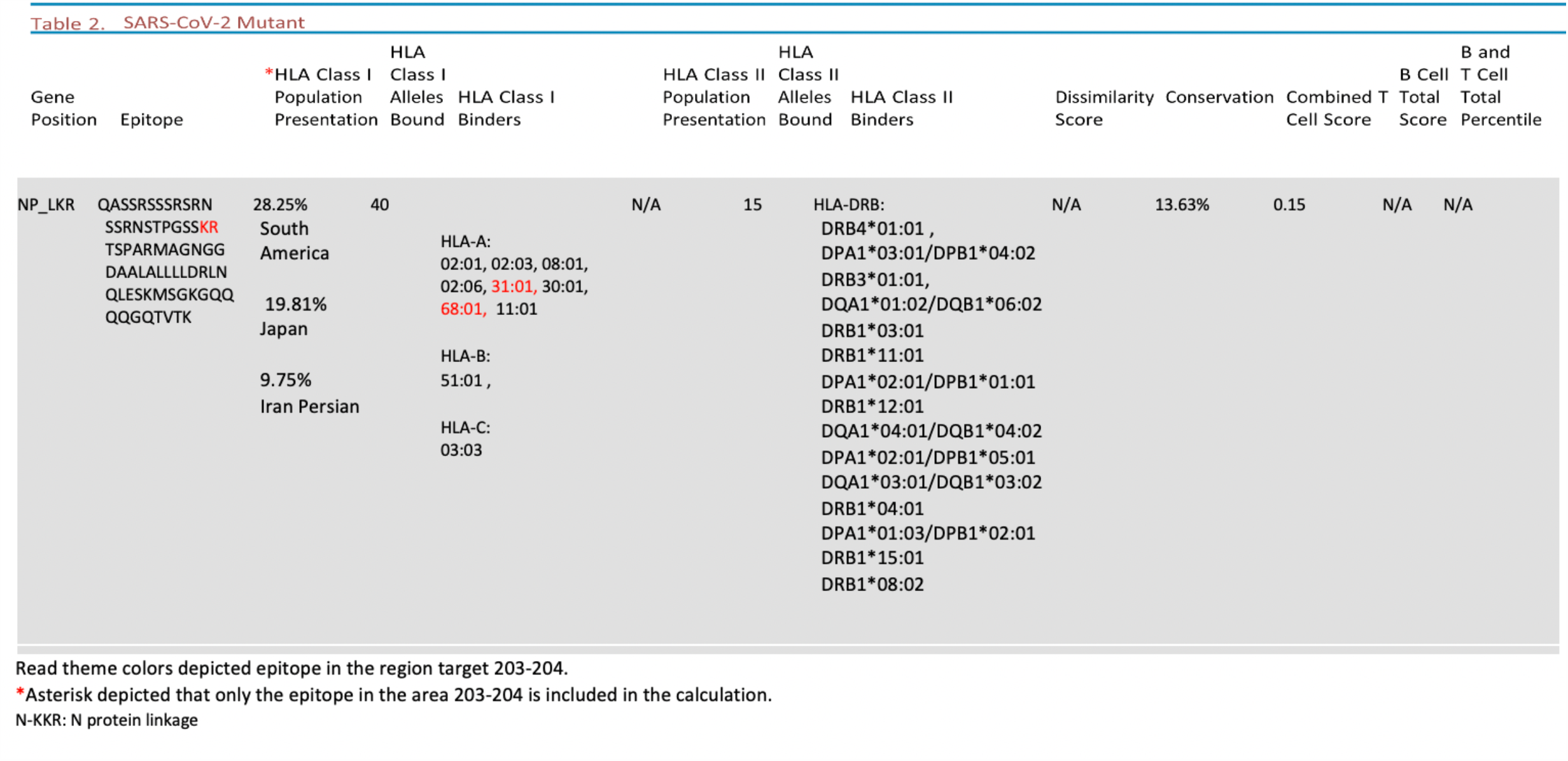
SARS-CoV-2 mutant G204R type N protein T cell HLA epitope predictions at gene position NP_LKR. HLA-class I binders depicted in red included only the epitope in the target region 203-204 in the calculation. Frequency of HLA class I representation within South American, Japanese, and Iranian populations is recorded as a frequency.

## 3. Discussion

Variations in Host Human Leukocyte Antigen (HLA) gene expression can influence antigenic presentation of coronavirus epitopes. The Nucleocapsid (N) proteins of many coronaviruses are highly immunogenic epitopes expressed abundantly during infection. Using genome sequencing and prediction models, we characterized a mutation in the N-protein which affected the viral structure, function, and immunogenicity.

In the case of SARS-CoV N protein, previous studies have demonstrated that there are three intrinsically disordered regions (IDRs) (residues 1–44, 182–247, and 366–422) which modulate the RNA-binding activity of the N-terminal domain (NTD) and the C-terminal domain (CTD) (11). The middle IDR, which we coined the linker region of SARS-CoV N protein (a.a. 182–247) LKR, and C-terminal IDR have both been implicated in the oligomerization of the N protein (12, 13). Five a.a variations at position 203 (R203K/M/S/I/G) within the SR-rich domain have been studied of the five variations, substitution R203K occurred most frequently in 68.09% of the mutated strains globally followed by the substitution G204R found in 67.94% mutated strains of the SARS-CoV-2 [19].

We identified phosphorylation sites in close proximity to the 203/204: RG>KR mutation and modeled the mutation’s potential role in the alteration of protein structure and favorable electrostatic interaction between positively charged Arg and the negatively charged P-Ser202/205. A decrease in structural instability could contribute to a potential increase in immunogenicity of the mutant variant.

The N protein wild type is known to be a representative antigen for the T-cell response in a vaccine setting, inducing SARS-specific T-cell proliferation and cytotoxic activity (14, 15). We discovered RG>KR mutated SARS-CoV-2 expands the ability for differential expression of the N protein epitope on Major Histocompatibility Complexes (MHC) of varying Human Leukocyte Antigen (HLA) origin. Specifically, the N protein LKR region K203 R204 of wild type (SARS-CoVs) and (SARS-CoV-2) observed HLA-A*30:01 and HLA-A*30:21, but mutant SARS-CoV-2 observed HLA-A*31:01 and HLA-A*68:01. Expression of HLA-A genotypes associated with the mutant strain occurred more frequently in all populations studied (South American, Japanese, and Iranian) with the most significant increase in frequency seen in the South Americans.

Major MHC-I molecules play a key role in the recognition of intracellular pathogens. HLA-A*31:01, associated with mutant and wild type strains is reported to be linked with carbamazepine (CBZ)-induced severe cutaneous adverse reactions (SCAR), including medicine reaction with eosinophilia and systemic symptoms (DRESS), Stevens-Johnson syndrome (SJS), and toxic epidermal necrolysis (TEN) (16). HL-A*68:01 was only associated with the mutant strain in our analysis. Two previous independent studies linked the HLA-A*68:01 allele, which is expressed at 5.2–25% allele frequency, with severe influenza disease during the 2009 influenza pandemic (17, 18).

The frequency of the HLA-A*68:01 allomorph is high among the indigenous populations globally including Southern America (http://www.allelefrequencies.net) and Australia (19, 20). HLA-A*68:01 is at low levels in most of SE Asia, particularly the indigenous populations reflected by low levels along the West Pacific Rim including Japan (19). If HLA-A*68:01 expression correlates with the manifestation of a more severe illness course, HLA-A*68:01 allele frequencies worldwide could positively correlate with the severity of coronavirus cases and a high number of deaths seen in south American countries like Brazil (21). On the other hand, low HLA-A*68:01 expression could correlate with the low number of COVID-19-attributable deaths seen in Japan as compared to other industrialized countries (22).

## 4. Conclusion

Findings in this study demonstrate how variations in Host Human Leukocyte Antigen (HLA) gene expression can influence the antigenic presentation of coronavirus epitopes. We identified phosphorylation sites in close proximity to the 203/204: RG>KR mutation and modeled the mutation’s potential role in the alteration of protein structure and favorable electrostatic interaction between positively charged Arg and the negatively charged P-Ser202/205. Major MHC-I molecules play a key role in the recognition of intracellular pathogens. Low HLA-A*68:01 expression could correlate with the low number of COVID-19-attributable deaths seen in Japan as compared to other industrialized countries. Importantly, we found that RG>KR mutated SARS-CoV-2 expands the ability for differential expression of the N protein epitope on Major Histocompatibility Complexes (MHC) of varying Human Leukocyte Antigen (HLA) origin.

## 5. Methods

### 5.1. Data retrieval and sequence alignment

We used BLASTP programs from the NCBI database search (23) to find the LKR N-protein sequence (43 nucleotides long) of all SARS-CoV-2. Conserved and varied residues were identified by using the WebLogo program (24-26). Multiple alignments were performed between full-length N-protein sequences on the EMBL-EBI server. Clustal Omega is used to apply mBed algorithms for guide trees. ClustalW alignment tools executed to output alignment format (27). We analyzed all available sequences available up to September 07th, 2020.

### 5.2. Structure modeling

The atomic coordinates of the N-terminal domain (NTD) and C-terminal domain (CTD) were obtained from the structure that is available in a Protein Data Bank (http://www.rcsb.org/pdb) (PDB ID: 6M3M, 6WZO)(28, 29). The tertiary structure of the full 419 a.a. sequencing of N-proteins was predicted using the IntFOLD5 server (PYMOL).

Sequences from residues 1-419 for N-protein native Sequence ID: YP_009724397.2 and mutant sequence ID: QIQ08827.1 were used in this study. All of the structures were visualized using PYMOL Chimera software Version 1.7.4 (30). The calculation procedure was almost the same as that in our previous works (31-34).

### 5.3. Model Validation

ProSA was used to measure the energy distribution of the N-protein structure (35, 36).

### 5.4. Identification of B-cell epitopes

In this subsection, we used the Immune Epitope Database (IEDB) (36) to determine linear B-cell epitopes operating the incorporated Chou & Fasman Beta-Turn prediction module (37). We supplied the FASTA sequence of the targeted protein as an input considering all default parameters. We also used the Discotope 2.0 (38) method to predict epitopes based on N-protein conformation and residue exposure.

### 5.5. Identification of T-cell epitopes

We used the TepiTool, a T cell Epitope Tool that is used for MHC class I and II binding predictions. The IEDB team’s recommendations were selected as defaults to automatically select the top peptides (39). In the MHC-I Binding Prediction feature, the default value provided is 1.0, i.e. all peptides with percentile rank ≤ 1.0 will be selected as predicted peptides. The list of representative alleles from different HLA supertypes was selected by the panel of 27 allele reference sets. The peptide selection criterion in this approach will always be predicted percentile rank. MHC-II binding prediction Results from the default value provided is 10.0, i.e. all peptides with percentile rank ≤ 10.0 will be selected as predicted peptides by using the panel of 26 most frequent alleles.

### 5.6. Prediction of protein subcellular localization using deep learning

The subcellular localization of wild and mutant N proteins was predicted using DeepLoc-1.0 software (40). Using the 419 sequence number at residues 203/204, a.a. of wild/ mutant strains were used to provide information about the subcellular localization of eukaryotic proteins using Neural Networks algorithm trained.

